# Amyloid-β oligomers are captured by the DNAJB6 chaperone: Direct detection of interactions that can prevent primary nucleation

**DOI:** 10.1101/2020.03.16.993790

**Authors:** Nicklas Österlund, Martin Lundquist, Leopold L. Ilag, Astrid Gräslund, Cecilia Emanuelsson

## Abstract

A human molecular chaperone protein, DNAJB6, is an efficient inhibitor of amyloid aggregation owing to a unique motif with conserved S/T-residues with high capacity for hydrogen bonding. Global analysis of kinetics data previously showed that especially the primary nucleation rate is inhibited. It was concluded that DNAJB6 achieves this remarkably effective and sub-stoichiometric inhibition by interacting not with the monomeric unfolded conformations of the amyloid-β (Aβ) peptide but with aggregated species. The pre-nucleation oligomeric aggregates are transient and difficult to study experimentally. Here we employed an approach to directly detect oligomeric forms of Aβ formed in solution by subsequent analysis with native mass spectrometry (native MS). Results show that the signals from the various forms of Aβ (1-40) oligomers were reduced considerably in the presence of DNAJB6, but not with a mutational variant of DNAJB6 in which the S/T-residues were substituted. With focus on DNAJB6 we could also detect signals that appear to represent DNAJB6 dimers and trimers to which varying amounts of Aβ is bound. These data provide direct experimental evidence that it is the oligomeric forms of Aβ that are captured by DNAJB6 in a manner which is dependent on the S/T residues. Strong binding of Aβ oligomers to DNAJB6 should indeed inhibit the formation of amyloid nuclei, in agreement with the previously observed decrease in primary nucleation rate.

## Introduction

Amyloid fibril formation by misfolded or intrinsically disordered proteins has recently been successfully described by kinetic models based on microscopic rate constants for fibril nucleation, fragmentation and elongation (1, 2). Nucleation can be divided into events which are only dependent on the monomer concentration (primary nucleation) and events which are dependent on both the monomer and fibril concentration (secondary nucleation). Amyloid-β (Aβ) peptide, a disease-related amyloidogenic agent in Alzheimer’s disease, is an intrinsically disordered peptide of 39-43 amino acid residues, which is very aggregation-prone. The two most abundant forms are the 40 and 42 residues long peptides, Aβ(1-40) and Aβ(1-42), with Aβ(1-40) being the most abundant and Aβ(1-42) with two additional hydrophobic residues being the more aggregation-prone and disease-related form (3). The Aβ amyloid aggregation process has been found by such kinetic analysis to be dominated by fibril catalyzed secondary nucleation (4). The difference in aggregation rates between Aβ(1-40) and Aβ(1-42) has also been shown to be due to a lower nucleation rate for Aβ(1-40), particularly the primary nucleation rate (5). Kinetic analysis gives insight into the different assembly rates underlying the formation of aggregates but does not include any detailed structure of the states along the aggregation pathway. The exact pathway for structural assembly of Aβ is currently not known in detail but a brief general overview is given here and summarized in Fig. 1.

**Figure 1.**
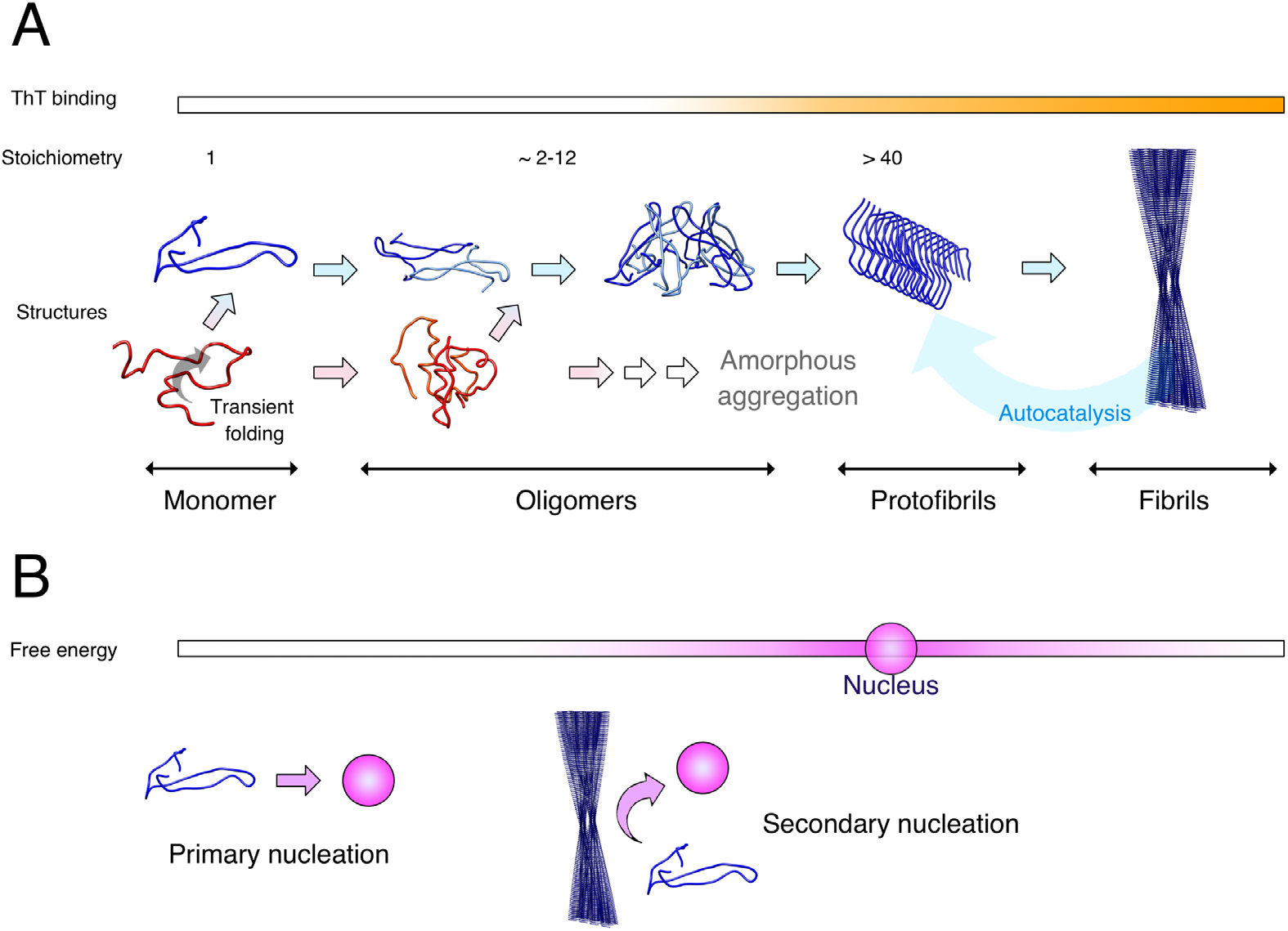
Overview of the Aβ aggregation process. **(A)** The Aβ peptide monomer is mostly unstructured (red) in solution but can exhibit a partial folding into a transient β-hairpin conformation (blue). Oligomerization proceeds through both unstructured and structured states. Oligomers are smaller and less well-organized assemblies with lower growth rate than protofibrils. Protofibrils eventually form large mature fibrils which catalyze the aggregation process in a positive feedback-loop (Autocatalysis). **(B)** Outline of the free energy landscape in which the pathway outlined in (A) proceeds. The darker color represents higher value of the free energy and the state with maximum free energy is termed “nucleus”, a state where addition of peptide monomers is more favorable than dissociation. Formation of nuclei can occur either through primary nucleation by association of pre-nucleation species (monomers, oligomers), or by secondary nucleation which is dependent on both pre-nucleation species and fibrils. Oligomers are here considered to be prenucleation species as they are intrinsically unstable and rapidly dissociate into monomers.

Monomeric peptides of Aβ are predominantly unstructured in solution, but can access a transient β-hairpin fold, which may be important for the aggregation process (6–9). The smallest peptide aggregates are termed oligomers, several definitions for these species are, however, used in the field based on, for example, size, growth rate, structure or function (10), and many studies use an operational definition based on what the employed method can detect. Studies have reported that various types of Aβ oligomers are toxic species formed in the amyloid assembly pathway (11–14). We here use an operational definition of oligomers (oligo – “a few”) as soluble assemblies of 2-12 peptides (9-60 kDa) that are detectable by native mass spectrometry (15). Larger oligomeric structures which have grown more fibril-like with an elongated linear shape are usually termed protofibrils (16, 17). Protofibrils can also be defined as the smallest Aβ structures that bind Thioflavin T (ThT), an amyloid specific dye which increases its fluorescence quantum yield upon binding to amyloid structures (18). It has been found by Fluorescence Correlation Spectroscopy that the smallest ThT active aggregate consists of around 60 Aβ peptides (19). These protofibrils then eventually form large fibrils. The term “nucleus” is defined as the smallest aggregate for which addition of a monomer is energetically more favorable than loss of a monomer unit (10). The nucleus therefore corresponds to the aggregated species with the highest surface energy (Fig 1B). The exact molecular details of Aβ amyloid nuclei are not known in detail.

The term oligomer is used in the literature to describe species both smaller and larger than the nucleus. A recent quantitative analysis of small on-pathway oligomers of Aβ(1-40) and Aβ(1-42) reveals that these oligomers dissociate more quickly than they convert to fibrillar species (20). Thus, monomers undergo multiple oligomerization events on the path to fibrils and the oligomers are highly transient and dynamic species. The small oligomers which are the topic of our present study are therefore to be considered as pre-nucleation species.

The link between changes in kinetic parameters for amyloid formation by Aβ and changes in directly observable structural peptide states of Aβ is not straightforward. Microscopy techniques can be used to monitor if formation and morphology of large fibrils correlate with changes observed from experimental kinetic assays. The smaller oligomeric aggregates are harder to study directly due to their transient and heterogenous nature, as well as their coexistence with monomers and large aggregates. The challenge is illustrated by the fact that the total Aβ(1-42) oligomer population, of various oligomerization states (21), has been found to reach at maximum 1.5% of the total monomer concentration (4). This represents a major experimental challenge as most biophysical techniques only report on an average of the monitored ensemble of states as weighted by their populations. One of the few available experimental techniques for direct detection of oligomers is native mass spectrometry (MS), where oligomeric peptide states can be observed individually and in parallel. We have previously used such an approach to describe the exact oligomeric states of Aβ(1-40) and Aβ(1-42) peptides in micellar environments (22).

In the current study, the oligomeric forms of Aβ(1-40) peptide are investigated using native MS, and the effect of the human chaperone DNAJB6 on Aβ oligomerization is studied. The anti-amyloid function of DNAJB6 was found when screening the human chaperones for suppressors of polyQ (polyglutamine) peptide aggregation (23). Since then we have characterized DNAJB6 as a remarkably efficient suppressor also of Aβ amyloid aggregations (24, 25) The protective function of DNAJB6 observed *in vitro* appears to be highly relevant also *in vivo*, with evidence provided using cells and a mouse disease model that showed considerably delayed aggregation and disease onset (26). A crucial role for DNAJB6 is emphasized by its identification, more than two decades ago, as *Mrj* (<u>*m*</u>ammalian <u>*r*</u>elative to <u>D</u>na*J*) in gene trapping studies with mural embryonic stem cells where *Mrj* mutants died already at the embryonal stage (27).

Our data with kinetic analysis of Aβ aggregation (28) reveal that that DNAJB6 is able to inhibit the primary nucleation of amyloid formation by binding aggregated Aβ species in a process that is dependent on its conserved S/T-residues. Inhibition requires only sub-stoichiometric molar ratios of DNAJB6. At high concentrations the DNAJB6 chaperone forms large megadalton oligomers which are in equilibrium with dissociated subunits in a concentration-dependent manner. The anti-aggregation effect of DNAJB6, attributed to the binding of oligomeric rather than monomeric forms of Aβ (28), is here extended upon and using native MS we directly demonstrate the capturing of pre-nucleation Aβ oligomers by DNAJB6.

## Results

### DNAJB6 efficiently suppresses the primary nucleation of Aβ(1-40) during amyloid formation

To investigate the interactions with the Aβ oligomers we have used DNJB6 (DNAJB6 WT) and the mutational variant (DNAJB6 S/T18A) in which the functionally important serine and threonine (S/T) residues in DNAJB6 were substituted into alanine (Fig. 2A). These residues surround a peptide-binding cleft at the interface between two monomers (Fig. 2B), according to our structural model (29). The functionality of DNAJB6 measured as its capacity to suppress aggregation and fibril formation by Aβ(1-40) is shown as a delay in the time-dependent ThT fluorescence increase during Aβ(1-40) aggregation (Fig. 2C, grey trace). Amyloid aggregation is delayed in the presence of DNAJB6 and the lag time is increased compared to the control sample with Aβ(1-40) only (green trace), with no effect on the growth rate. This is typical for inhibition of primary nucleation and in agreement with previous results for DNAJB6 with Aβ(1-42) (24), and the low amounts sub-stoichiometric of DNAJB6 required was 10-fold lower for Aβ(1-40) (Fig. S1). Such delay in aggregation was neither observed for the mutational variant S/T18A of DNAJB6 (cyan trace), nor for crosslinked DNAJB6 (purple trace), which is locked in oligomeric states, as revealed by denaturing electrophoresis (Fig. S2), and dissociation of DNAJB6 oligomer is prevented.

**Figure 2.**
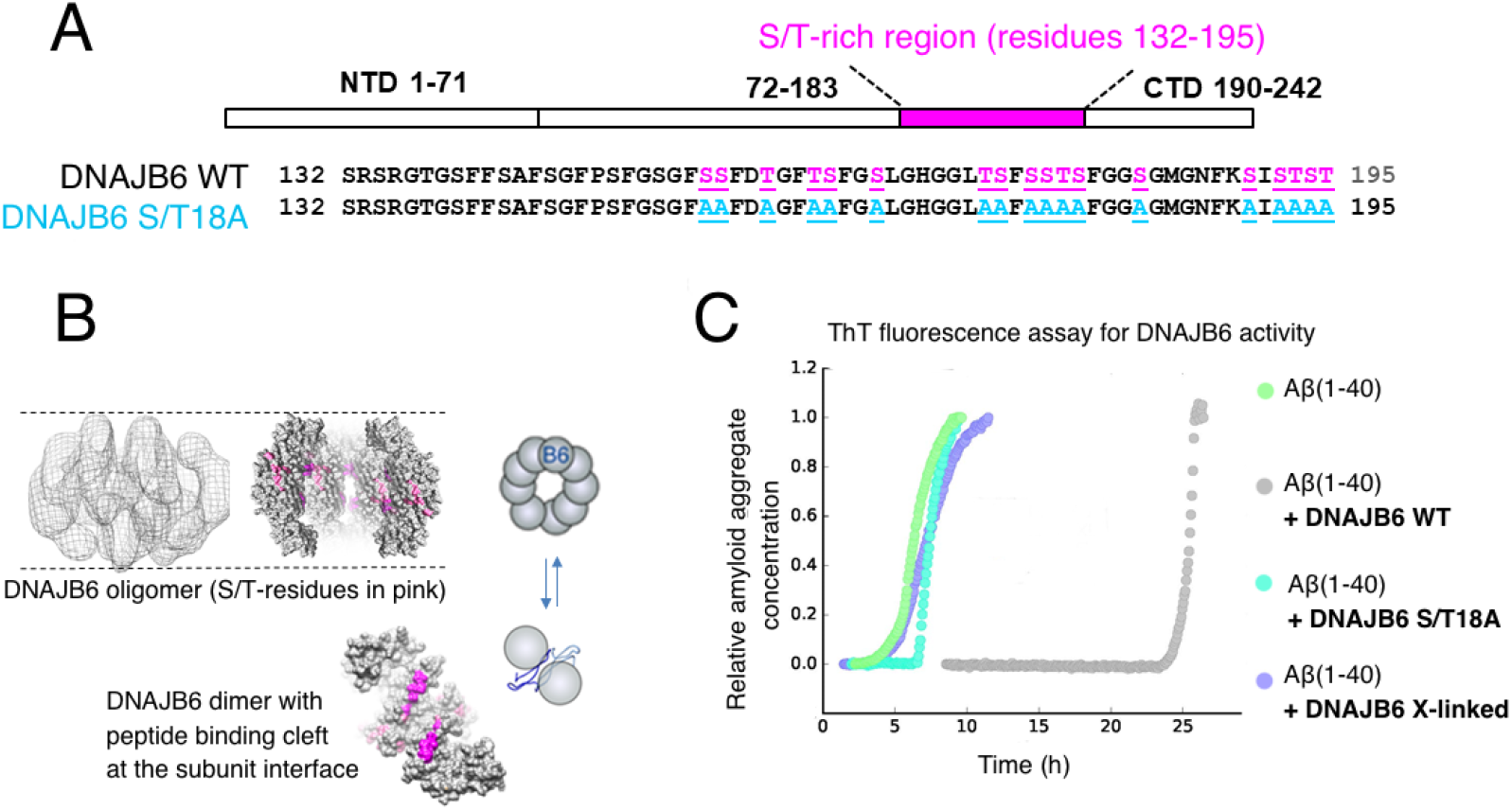
The chaperone DNAJB6 with functionally important ST-residues can suppress fibril formation by Aβ(1-40). (A) Conserved serine- and threonine- (S/T) residues in DNAJB6 are high-lighted in pink. 18 S/T-residues are substituted with alanine-residues in a mutational variant of DNAJB6 referred to as S/T18A. **(B)** Structural model of DNAJB6, with an outline in cartoon of DNAJB6 oligomers in equilibrium with dissociated subunits. The S/T-residues are proposed to bind Aβ in a peptide-binding cleft formed at the interface between two monomeric subunits. **(C)** The capacity of DNAJB6 to suppress fibril formation by Aβ(M1-40) determined by ThT fluorescence measurement. The concentration of DNAJB6 used here is 0.003 μM and the molar ratio of Aβ(1-40) to DNAJB6 is 1:0.0002. Color code: Aβ(M1-40) only (green), Aβ(M1-40) + DNAJB6 WT (grey), Aβ(M1-40) + DNAJB6 mutational variant ST18A (cyan), Aβ(M1-40) + crosslinked DNAJB6 WT (purple).

### Oligomers of Aβ(1-40) are detected by native MS

Positive ion mode native MS analysis of Aβ(1-40) in ammonium acetate solution pH 7 reveals an m/z distribution where the major peaks correspond to monomer ions with 2-5 positive charges and smaller amounts of dimers, trimers and tetramers are also detected in several different charge states (Fig. S3), in agreement with previous observations (15, 22).

The low relative intensity for oligomers detectable in native MS is in agreement with the conclusions based on a number of other methods that Aβ oligomers only constitute a few percent of the total Aβ peptide population (21).

Oligomers can overlap in the mass/charge (m/z) dimension of the mass spectrometer (e.g. a monomer 2+ ion will overlap with a dimer 4+ ion). We therefore annotate peaks by their oligomeric state/charge (n/z) ratio. Overlapping n/z states can often be deconvoluted using the ^13^C isotopic distribution (Fig. S4) or by using ion mobility measurements.

The signal intensities detected in native MS cannot directly be used to quantify the absolute solution state concentration of each species. It has to be considered that monomers and different oligomeric states may not have the same ionization efficiency and that oligomers may, to some extent, dissociate or associate in the gas phase. We have normalized all signals by taking relative intensities, defined as the ratio between the mass intensity of a particular ion signal and the sum of the mass intensity of all detected signals in a mass spectrum. We then consider the changes of relative intensity, within each specific charge state of a certain oligomer, in samples without or with a 1 h incubation in solution. These values we consider relevant as proxy reporter for the concentrations in solution.

### Pre-incubation with DNAJB6 decreases the amount of free Aβ(1-40) oligomers

Aliquots of Aβ(1-40) were pre-incubated in solution (37°C, 1h), in the absence or presence of DNAJB6 (WT or the mutational variant S/T18A), to permit formation of oligomers as outlined in Fig 3A. The control samples pre-incubated in absence of DNAJB6 were supplied with a corresponding amount of DNAJB6 right before native MS, to avoid differences in Aβ(1-40) ionization efficiency caused by addition of DNAJB6. Aβ(1-40) peaks were normalized relative to the sum of all Aβ(1-40) signals in the mass spectrum.

**Figure 3.**
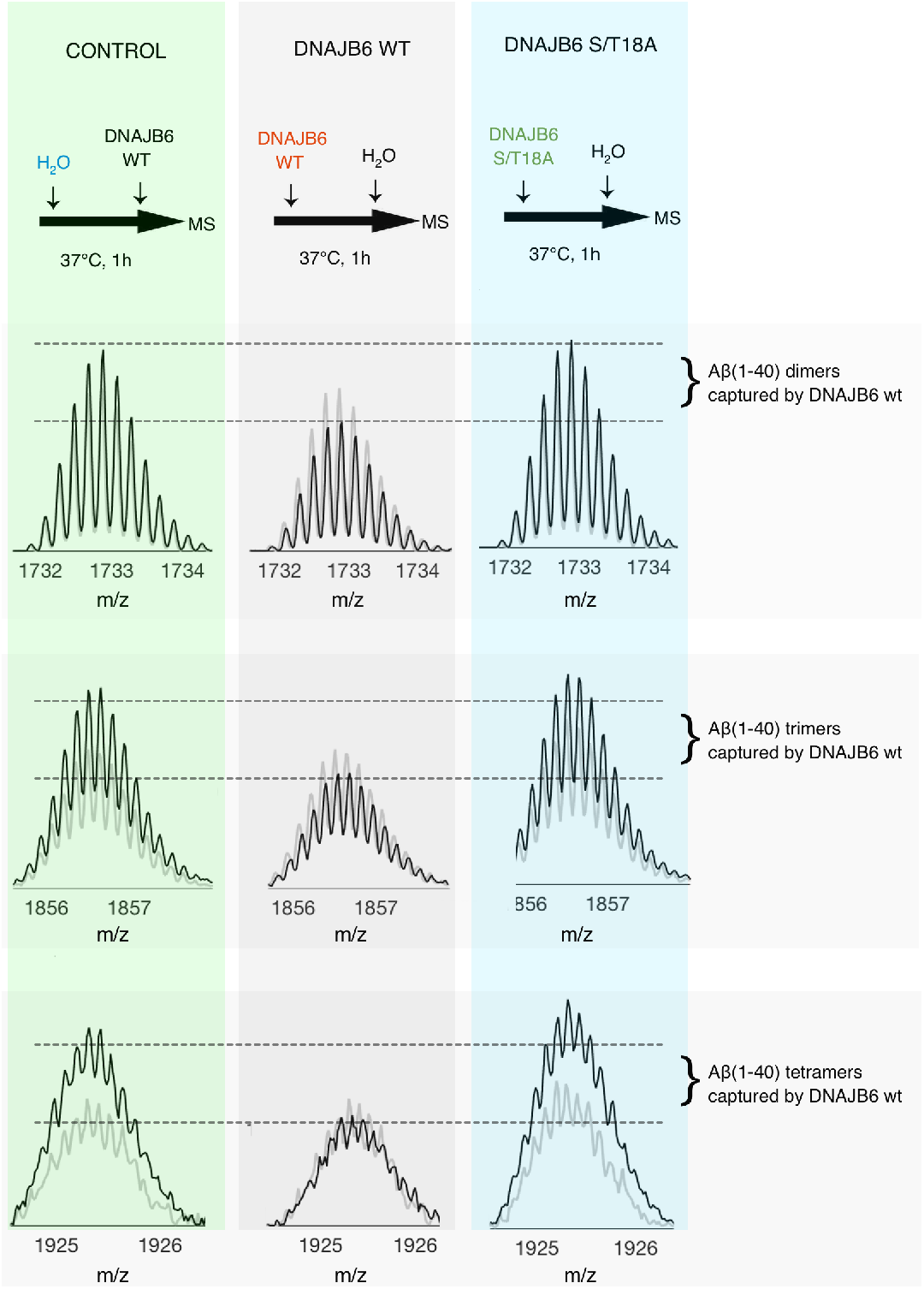
Aβ(1-40) oligomers are captured by DNAJB6. **(A)**, an outline of how the samples were pre-incubated for possible formation of Aβ(1-40) oligomers in solution and then injected for detection of Aβ oligomers in the gas phase by native MS. The control sample was supplied with DNAJB6 protein just before the IM-MS analysis to account for changes in ionization efficiency by added DNAJB6. **(B), (C)** and **(D)**, signals from Aβ40 dimers (n/z=2/5), trimers (n/z=3/7) and tetramers (n/z=4/9), respectively. The signals at time = 0 are shown as shaded lines and at time 1 h as black lines. Signals are normalized against the corresponding signal from the Aβ(1-40) monomer. A freshly prepared preparation of Aβ(1-40) was diluted in 10 mM ammonium acetate pH 7 solution to a final peptide concentration of 10 μM and incubated 1 h at 37°C, either without DNAJB6 (control, green), or in the presence of DNAJB6 WT (grey) or DNAJB6 S/T18A (cyan), the mutational variant with 18 S/T to A substitutions, at a molar ratio of Aβ(1-40) to DNAJB6 of 1:0.1.

The n/z signals 2/5, 3/7 and 4/9 represent relatively high intensity signals for the dimeric, trimeric and tetrameric states of Aβ(1-40) respectively, which do not overlap with other oligomeric signals in the m/z dimension as they have odd number of charges (Fig S3B). These n/z states were monitored as presented in Fig. 3B-D. The signal intensities for Aβ(1-40) oligomers increased after 1 h (left panels, black lines) compared to time zero (left panels, shaded lines), when pre-incubated in the absence of DNAJB6. This increase reports on the amount of Aβ(1-40) oligomers formed in solution during 1 h. In contrast, the signal intensities did not increase after pre-incubation in presence of DNAJB6 (middle panel), instead there was a decrease. This indicates that that the free dimers, trimers and tetramers of Aβ(1-40) have been captured and removed from the soluble peptide pool by DNAJB6 during the pre-incubation. This is not observed upon pre-incubation with the mutational variant of DNAJB6 which lacks the S/T residues (right panel).

Experiments were also conducted to evaluate the concentration dependency of the DNAJB6 effect. As shown in Fig. 4 for the signals from Aβ(1-40) with n/z = 2/5, the decrease is linear with the DNAJB6 concentration (inset). The same trend is observed also for the other oligomeric states (Fig. S5). This illustrates that a larger amount of DNAJB6 is able to capture a larger amount of Aβ oligomers.

**Figure 4.**
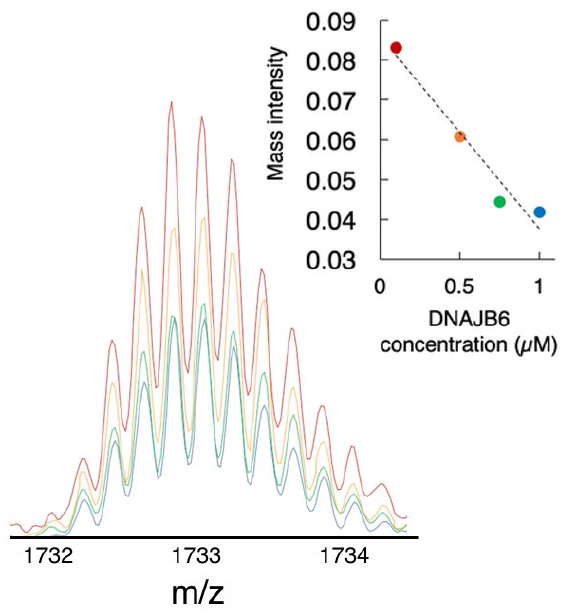
Concentration-dependence of the decreased signal from Aβ(1-40) dimer after pre-incubation with DNAJB6. The experiment was performed as described in Fig. 3 using 10 μM Aβ(1-40) and 0.1-1 μM DNAJB6 WT and the image shows the signal from Aβ (1-40) dimer (n/z=2/5). The insert shows that the decrease in signal intensity is linear with respect to DNAJB6 concentration.

The data in Fig. 3 and Fig. 4 show that our native MS data provide direct experimental observations supporting the conclusion from kinetic analyses that DNAJB6 can remove the oligomeric forms of Aβ from solution, thereby preventing the growth and the proliferation of Aβ aggregates. The weak effect of the mutational variant DNAJB6 S/T18A confirms the importance of the S/T-residues for interaction between the DNAJB6 chaperone and the oligomeric pre-nucleation Aβ aggregates.

### Changes in Aβ(1-40) oligomer intensities are dependent on oligomeric state and charge state

The observations for the three n/z signals shown in Fig. 3 are observed also for the other charge states, but to a varying extent (Fig. S6-S8). In the following we consider the change of relative intensity, within each oligomer and specific charge state, as a proxy reporter for the solution concentration of that species. The relative intensity I_R_ is defined as the ratio between the mass intensity of a particular ion signal and the sum of the mass intensity of all detected Aβ signals in the mass spectrum. The change in relative intensity (I_R(1 h)_/I_R (0 h)_), is the relative intensity at the end compared to the start of the 1 h pre-incubation in solution.

The change in relative intensity is shown for each detected Aβ ion in Fig. 5. Only n/z signals where the exact oligomeric state could be distinguished in mass dimension by the ^13^C isotopic pattern (Fig. S4) were evaluated. The change in relative intensity over 1h of pre-incubation is shown for each charge state and each oligomer state of Aβ(1-40) without DNAJB6 (Fig. 5A, left panel, green), with DNAJB6 (Fig. 5A, middle panel, grey) and with the DNAJB6 S/T18A mutant (Fig. 5A, right panel, blue). The effect per oligomeric state when averaging over all charge states is shown in Fig. 5B. (without DNAJB6, with DNAJB6 WT and with DNAJB6 S/T18A). Some observations can be made of the events during the 1 h pre-incubation:

i. Without DNAJB6 all Aβ(1-40) oligomeric species increase in relative intensity. Higher oligomeric states increase more than lower oligomeric states. (e.g. tetramers more than dimers, Fig. 5A, left panel).
ii. The relative intensity of low charged n/z signals increase more than high charged n/z signals, and even monomers show a shift towards lower charge states upon incubation (Fig. 5A, left panel). Charging in electrospray ionization under native conditions is generally proportional to the solvent accessible surface area of the protein (30). This charge-state dependence thus indicates that during the incubation there is an increase of low charge/compact forms of the Aβ oligomers.
iii. DNAJB6 stops formation of new oligomers and removes already formed oligomers during the 1 h pre-incubation (Fig. 5A, middle panel). The removal of oligomers is most pronounced for the low charge/compact forms of the Aβ oligomers.
iv. DNAJB6 mutant ST18A does not remove already formed oligomers but to some extent hinders formation of new oligomers (Fig. 5A, right panel). The molar ratio of Aβ to DNAJB6 is high (1:0.1) compared to that required for inhibition of aggregation (Fig. 2) so unspecific effects may occur.

**Figure 5.**
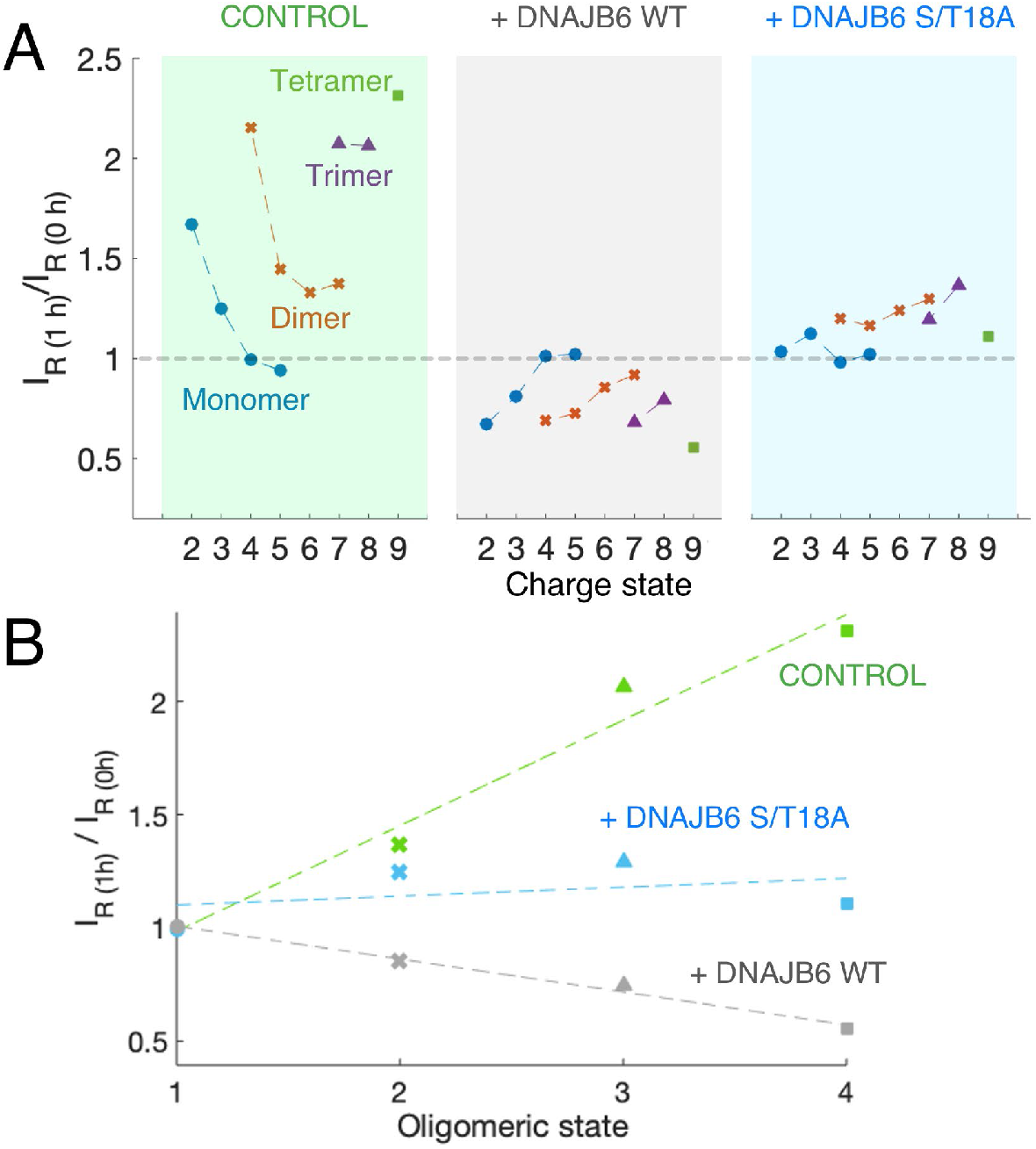
Change in relative intensity for detected mass spectrometry signals of various forms of Aβ(1-40) following one hour of incubation in solution. Signals are normalized against the total sum of Aβ(1-40) signals. **(A)** Aβ(1-40) samples were injected before or after 1 h pre-incubation in solution, either without DNAJB6 (left/green), or with wild-type DNAJB6 (middle/gray) or with the S/T18A mutant of DNAJB6 (right/cyan). Signals are shown for each oligomeric state and each charge state from the mass spectra shown in Fig. S4, for n/z signals where the exact oligomeric stated could be distinguished in mass dimension by the 13C isotopic pattern. **(B)** The effect per oligomeric state, when averaging over all charge states.

The ion mobility of the Aβ ions was measured to detect possible shifts in the Aβ conformational ensemble, as suggested by the charge state distribution analysis. Although shifts in the drift time profiles of n/z signals were observed there were no clear trends for shifts towards compact or extended states upon incubation with DNAJB6 WT (Fig. S9). It should be noted that ion mobility reports on gas phase structure rather than solution state structure. The difference in structure in the two different environments will be especially large for weakly structured proteins such as Aβ, as the apolar vacuum of the mass spectrometer may stabilize structure due to increased intramolecular hydrogen bonding (31, 32). The analysis of charge-state distributions should therefore be a better tool in this case, as this reports on structure changes in the solution state where the ionization process takes place.

### Aβ(1-40) binds to dissociated DNAJB6 oligomers

A higher concentration (37 μM) of DNAJB6 was injected into the mass spectrometer for direct detection of the chaperone itself. DNAJB6 monomer signals, with a narrow distribution of two major charge states of 9+ and 10+, were detected (Fig. 6A). The narrow charge-state distribution and low charge is indicative of a folded state. Well-established theory for charging of folded proteins predict that a 27 kDa protein (DNAJB6 monomer), would carry an average charge of 10.5 (33, 34). No other DNAJB6 signals were observed. Previous studies show that DNAJB6 occurs as oligomers in the MDa range (29), in equilibrium with dissociated subunits. Such large oligomers are beyond the mass range of the here used mass spectrometer and therefore not possible to observe.

**Figure 6.**
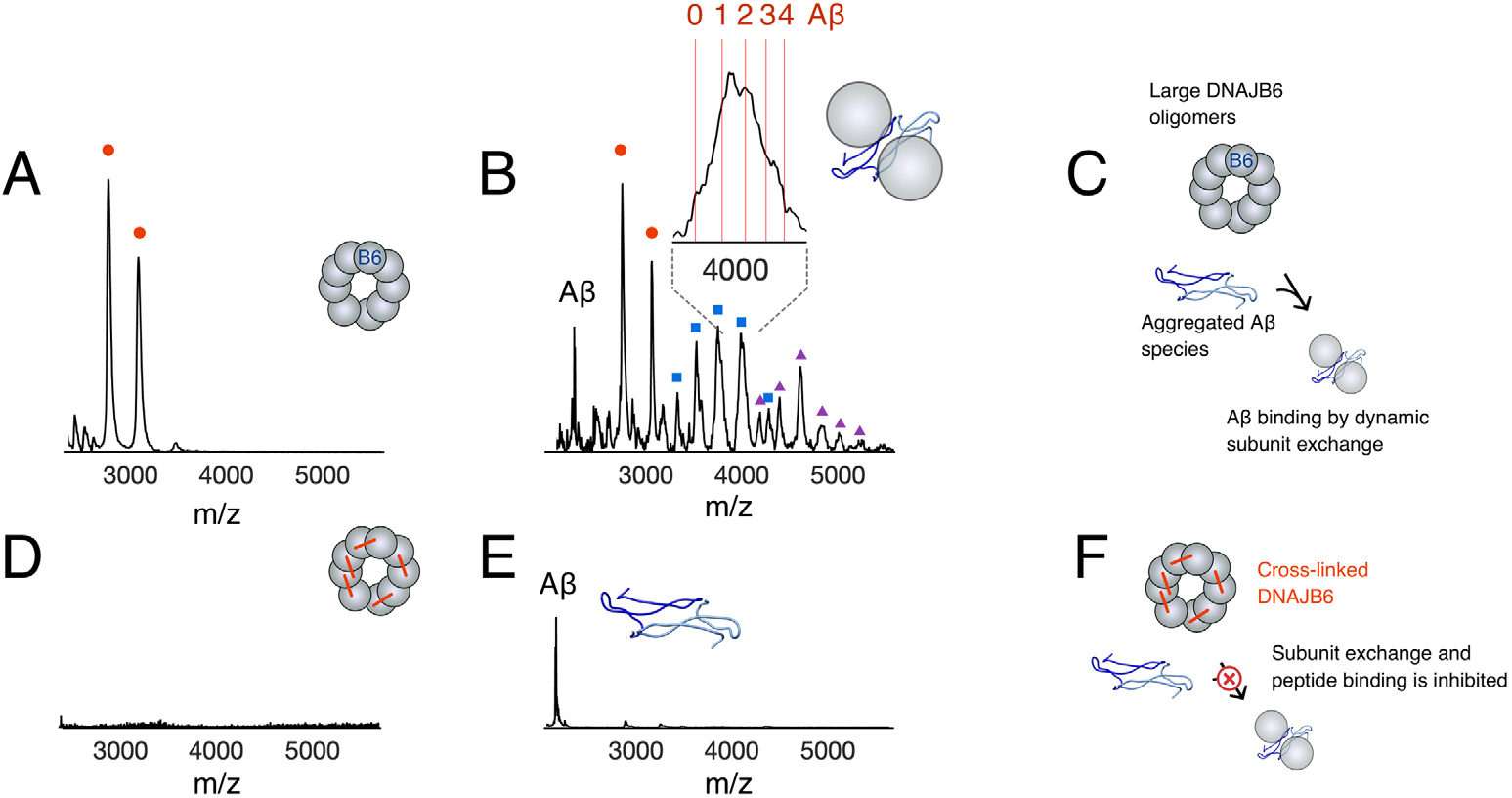
DNAJB6 oligomer dissociation and binding of Aβ(1-40) in small Aβ(1-40)-DNAJB6 complexes. **(A)** Native MS spectrum of 37 μM DNAJB6. The red circles indicate signals (+9/+10) for folded DNAJB6 monomers. **(B)** Native MS spectrum of 37 μM DNAJB6 after pre-incubation with 1 μM Aβ(1-40). Signals appear that correspond to DNAJB6 dimers (blue squares) and trimers (purple triangles). Closer inspection of the peaks shows that the DNAJB6 peaks are shifted towards masses indicating binding of 1-4 Aβ peptides (insert). **(C)** Proposed binding of Aβ oligomers to DNAJB6 oligomers following oligomer dissociation and binding of Aβ to the S/T-rich region in a complex that hinders further aggregation into amyloid nuclei and primary nucleation, as observed in analyses of aggregation kinetics, and suppression of amyloid fibril formation (Fig. 2C). **(D)** Native MS spectrum of 37 μM cross-linked DNAJB6. No signals corresponding to monomeric DNAJB6 are detected. **(E)** Native MS spectrum of 37 μM cross-linked DNAJB6 after pre-incubation with 1 μM Aβ(1-40). Only peaks from Aβ are detected. **(F)** As in (C), but following cross-linking of DNAJB6 that inhibits oligomer dissociation, there is no suppression of amyloid fibril formation (Fig. 2C) and no binding of Aβ-oligomers.

Interestingly, new peaks appeared when DNAJB6 (37 μM) was supplied with Aβ(1-40) (1 μM) immediately prior to injection (Fig. 6B). These new peaks show peak broadening and mass shifts to masses slightly higher than twice the mass of the observed DNAJB6 monomer peaks. This could correspond to DNAJB6 dimers and trimers, with masses in agreement with 1-4 copies of the Aβ(1-40) peptides bound, as shown in the inset in Fig. 6B. This suggests that dissociation of the DNAJB6 oligomers occurs in presence of Aβ(1-40), and that small complexes form between dissociated DNAJB6 and captured Aβ(1-40), as outlined in Fig. 6C.

However, if DNAJB6 was crosslinked, no peaks corresponding to DNAJB6 monomer signals were detected upon injection (Fig. 6D) and no new peaks representing small complexes of dissociated DNAJB6 and captured oligomers were detected in presence of Aβ(1-40) (Fig. 6E), only signals for free Aβ(1-40) could be observed. Crosslinking of DNAJB6 prevents oligomer dissociation (Fig. S2), and therefore no binding of (Aβ(1-40) occurs as outlined in Fig. 6F, which explains that crosslinked DNAJB6 cannot suppress fibril formation (Fig. 2C).

## Discussion

In this study we have detected small Aβ oligomers using native mass spectrometry, enabling direct observation of individual oligomeric states during the Aβ aggregation process. The changes in these directly observable molecular states agree well with changes in amyloid formation kinetics upon modulation of the primary nucleation rate for Aβ by the human chaperone protein DNAJB6.

Our herein presented ThT kinetic data on Aβ(1-40) and DNAJB6 (Fig. 2) are in line with previous data showing that DNAJB6 is remarkably efficient in suppressing fibril formation of Aβ(1-42) at very low (1:0.01) molar ratio of peptide to chaperone (28). It should be noted that the molar ratios of DNAJB6 to Aβ(1-40) used here to successfully suppress Aβ amyloid formation are even lower (at least ten-fold lower, i.e. the molar ratio of DNAJB6 to Aβ(1-40) is in the order of 1:0.001). This is in agreement with the finding that the primary nucleation rate is intrinsically lower for Aβ(1-40) as compared to Aβ(1-42) (5) and that oligomerization is less extensive for Aβ(1-40) as compared to Aβ(1-42) (15, 22). Less DNAJB6 is consequently needed to capture these very few formed oligomeric aggregates.

The current model for *in vitro* inhibition of Aβ aggregation by DNAJB6 is that the chaperone binds the oligomeric Aβ aggregates strongly and removes them from the soluble Aβ pool available for amyloid aggregation (24). Thereby the formation of primary Aβ nuclei is prevented or delayed. This also means that the concentration of active DNAJB6 in the solution decreases over time, with loss of inhibitory effect as a result. The depletion of Aβ oligomers over time (Fig. 3-5) as well as the formation of Aβ-DNAJB6 complexes (Fig. 6B) are directly observed here, giving independent support for this model.

The capture of the Aβ oligomers and their removal from solution, which does not occur to the same extent in the DNAJB6 ST18A-mutant, is obviously dependent on S/T-residues in the chaperone, which we believe form intermolecular hydrogen bonds to Aβ. This is reminiscent of how Aβ peptides are captured by Z_Aβ3_-related affibodies (35, 36). In such systems hydrogen bonds are formed between the affibody and the Aβ backbone, forming a complex where a monomeric Aβ peptide is captured in its β-hairpin state. DNAJB6 could similarly be imagined to bind oligomeric forms of β-hairpin Aβ via hydrogen bonding, as oligomers undergo a process of reorganization driven by interchain hydrogen bonding interactions (37). DNAJB6 might bind and stabilize oligomeric β-hairpin Aβ with high surface energies by hydrogen bonding to hydroxyl groups of the S/T-rich region in DNAJB6. Interestingly, recent data show that transthyretin is also, as DNAJB6, very efficient in preventing Aβ aggregation by inhibiting the primary nucleation (38). It has also been shown that a hydrogen-bond network is important for structural stability of transthyretin (39).

It is interesting to note that low charged/compact forms of Aβ oligomers, which increased most during 1 h pre-incubation in solution (Fig. 5A, left panel), were most efficiently captured by DNAJB6 (Fig. 5A, middle panel). It is intriguing to speculate that these charge states report on the most compact β-hairpin forms of Aβ in solution (Fig 1A), as solution state species with smaller surface area generally produce ions of lower charge in electrospray ionization. Charge state distribution analysis has previously been used to study the unfolded ensemble of intrinsically disordered proteins (40). The change in electrospray charging for the disordered amyloidogenic protein α-synuclein has for example been studied upon changes in solvent, pH and upon binding to ligands (41, 42). A similar systematic study of how the Aβ charge state distribution changes upon modulation of conditions is not presently available.

The strongly bound Aβ-DNAJB6 complexes observed *in vitro* are most likely not as long lived *in vivo,* where other downstream processes would be present. Currently, we can only speculate what the fate of the Aβ species captured by DNAJB6 can be under cellular conditions. Presumably the bound Aβ can be released and sent for proteasomal degradation in a cycle involving components such as Hsp70 and ATP (26).

In conclusion, we demonstrate in this study that the amount of Aβ oligomers detectable by native MS is considerably lowered if Aβ monomers are pre-incubated in the presence of DNAJB6 chaperone, a process which is dependent on the S/T-residues in DNAJB6. The effect of the chaperone is largest for larger Aβ oligomers and for low charged/compact forms of Aβ oligomers that may be the most compact β-hairpin forms. Detection of peaks corresponding to DNAJB6 dimers and trimers with mass shifts appear to represent a direct observation of Aβ oligomers captured by dissociated DNAJB6 subunits. This demonstrates the usability of native MS to study directly observable peptide states during Aβ peptide aggregation, as a complement to the information acquired from kinetic parameters.

## Experimental procedures

### Sample preparation

The amino acid numbering used here refers to the amino acid sequence of DNAJB6 isoform B (UniProt KB accession number O75190). Expression of DNAJB6 protein was performed at the Lund Protein Production Platform, Lund University (http://www.lu.se/lp3), as previously described (24, 28). Crosslinking of the DNAJB6 oligomers with the crosslinker BS3 which is specific for primary amines was performed at 50 μM concentration of DNAJB6 and a 3-fold molar excess of crosslinker to primary amines, as described previously (29). Prior to mass spectrometry. the buffer of the purified DNAJB6 was exchanged into 10 mM ammonium acetate pH 7 solution using Micro Biospin P6 centrifuge columns (Bio-Rad) and protein concentration determined using a NanoDrop spectrophotometer (Thermo Fisher Scientific, Stockholm, Sweden) and recombinant human Aβ(1-40) purchased from Alexotech AB (Umeå, Sweden) as lyophilized peptide was dissolved in 15% ammonium hydroxide, sonicated in an ice-water bath for 1 minute and diluted in 10 mM ammonium acetate pH 7.0 to a final peptide concentration of 10 μM.

### Activity measurements

The capacity of DNAJB6 proteins to suppress fibril formation by Aβ(1-40) was determined using ThT fluorescence as previously described (28). Recombinant human Aβ(M1-40) was expressed tag-free from a PetSac plasmid and purified as described previously (5, 43). Fresh monomer was isolated by size exclusion chromatography in 20 mM sodium phosphate, 0.2 mM EDTA, pH 7.4 just prior to setting up the kinetics experiments to remove any aggregated species. DNAJB6 was at a concentration of 0.003 μM, Aβ(1-40) at 10-28 μM and ThT 40 μM for detection of fibrils.

### Native IM-MS

A Waters Synapt G2S hybrid mass/ion mobility spectrometer equipped with a nano-electrospray source was used for analysis. Samples were injected using nano-electospray by commercial metal coated glass injectors (Thermo Scientific). Ionization was performed in positive ion mode and the instrument parameters were as follows: Capillary voltage 1.7 kV, Sampling cone 40 V, Source offset 80 V, Trap gas 10 mL/min, Helium Gas Flow 100 mL/min, IMS gas flow 50 mL/min, IMS wave velocity 750 m/s, IMS wave height 24 V. To detect signals from Aβ(1-40) a freshly prepared preparation of Aβ (1-40) was diluted in 10 mM ammonium acetate pH 7 solution to a final peptide concentration of 10 μM and injected without or with pre-incubation with DNAJB6 for 1 h at 37°C, at a molar ratio of Aβ (1-40) to DNAJB6 of 1:0.1 (i.e. concentration of DNAJB6 was 1 μM), or as otherwise stated. To the control samples, DNAJB6 was added after pre-incubation just before injection to avoid differences in ion suppression. To detect signals from DNAJB6 a higher concentration was used for injection (37 μM, 1 mg/ml)

## Acknowledgments

Professor Sara Linse is thanked for valuable discussions. This work was supported by the Crafoord Foundation (to CE), The Alzheimer Foundation (to CE), The Swedish Research Council (to AG), The Sven and Lilly Lawski Foundation (to NÖ), the Olle Engkvist Byggmästare Foundation (to LLI). The Native IM-MS infrastructure was supported by a SciLifeLab Technology Development Grant from the Faculty of Science at Stockholm University (to LLI). This work was facilitated by recombinant expression of DNAJB6 protein performed at the Lund Protein Production Platform, Lund University (http://www.lu.se/lp3).

## References

1. Meisl, G., Kirkegaard, J. B., Arosio, P., Michaels, T. C. T., Vendruscolo, M., Dobson, C. M., Linse, S., and Knowles, T. P. J. (2016) Molecular mechanisms of protein aggregation from global fitting of kinetic models. Nat. Protoc. 11, 252–72

2. Knowles, T. P. J., Waudby, C. A., Devlin, G. L., Cohen, S. I. A., Aguzzi, A., Vendruscolo, M., Terentjev, E. M., Welland, M. E., and Dobson, C. M. (2009) An analytical solution to the kinetics of breakable filament assembly. Science (80-.). 326, 1533–1537

3. Walsh, D. M., and Selkoe, D. J. (2007) Aβ oligomers - A decade of discovery. J. Neurochem. 101, 1172–1184

4. Cohen, S. I. A., Linse, S., Luheshi, L. M., Hellstrand, E., White, D. A., Rajah, L., Otzen, D. E., Vendruscolo, M., Dobson, C. M., and Knowles, T. P. J. (2013) Proliferation of amyloid-42 aggregates occurs through a secondary nucleation mechanism. Proc. Natl. Acad. Sci. 110, 9758–9763

5. Meisl, G., Yang, X., Hellstrand, E., Frohm, B., Kirkegaard, J. B., Cohen, S. I. A., Dobson, C. M., Linse, S., and Knowles, T. P. J. (2014) Differences in nucleation behavior underlie the contrasting aggregation kinetics of the Aβ40 and Aβ42 peptides. Proc. Natl. Acad. Sci. 111, 9384–9389

6. Abelein, A., Abrahams, J. P., Danielsson, J., Gräslund, A., Jarvet, J., Luo, J., Tiiman, A., and Wärmländer, S. K. T. S. (2014) The hairpin conformation of the amyloid β peptide is an important structural motif along the aggregation pathway. J. Biol. Inorg. Chem. 19, 623–34

7. Sandberg, A., Luheshi, L. M., Sollvander, S., Pereira de Barros, T., Macao, B., Knowles, T. P. J., Biverstal, H., Lendel, C., Ekholm-Petterson, F., Dubnovitsky, A., Lannfelt, L., Dobson, C. M., and Hard, T. (2010) Stabilization of neurotoxic Alzheimer amyloid- oligomers by protein engineering. Proc. Natl. Acad. Sci. 107, 15595–15600

8. Lendel, C., Bjerring, M., Dubnovitsky, A., Kelly, R. T., Filippov, A., Antzutkin, O. N., Nielsen, N. C., and Härd, T. (2014) A hexameric peptide barrel as building block of amyloid-β protofibrils. Angew. Chemie - Int. Ed. 53, 12756–12760

9. Kreutzer, A. G., and Nowick, J. S. (2018) Elucidating the Structures of Amyloid Oligomers with Macrocyclic β-Hairpin Peptides: Insights into Alzheimer’s Disease and Other Amyloid Diseases. Acc. Chem. Res. 51, 706–718

10. Arosio, P., Knowles, T. P. J., and Linse, S. (2015) On the lag phase in amyloid fibril formation. Phys. Chem. Chem. Phys. 17, 7606–7618

11. Meier, J. J., Kayed, R., Lin, C.-Y., Gurlo, T., Haataja, L., Jayasinghe, S., Langen, R., Glabe, C. G., and Butler, P. C. (2006) Inhibition of human IAPP fibril formation does not prevent beta-cell death: evidence for distinct actions of oligomers and fibrils of human IAPP. AJP Endocrinol. Metab. 291, E1317–E1324

12. Conway, K. A., Lee, S.-J., Rochet, J.-C., Ding, T. T., Williamson, R. E., and Lansbury, P. T. (2000) Acceleration of oligomerization, not fibrillization, is a shared property of both alpha-synuclein mutations linked to early-onset Parkinson’s disease: Implications for pathogenesis and therapy. Proc. Natl. Acad. Sci. 97, 571–576

13. Viola, K. L., and Klein, W. L. (2015) Amyloid β oligomers in Alzheimer’s disease pathogenesis, treatment, and diagnosis. Acta Neuropathol. 129, 183–206

14. De, S., Wirthensohn, D. C., Flagmeier, P., Hughes, C., Aprile, F. A., Ruggeri, F. S., Whiten, D. R., Emin, D., Xia, Z., Varela, J. A., Sormanni, P., Kundel, F., Knowles, T. P. J., Dobson, C. M., Bryant, C., Vendruscolo, M., and Klenerman, D. (2019) Different soluble aggregates of Aβ42 can give rise to cellular toxicity through different mechanisms. Nat. Commun. 10.1038/s41467-019-09477-3

15. Bernstein, S. L., Dupuis, N. F., Lazo, N. D., Wyttenbach, T., Condron, M. M., Bitan, G., Teplow, D. B., Shea, J. E., Ruotolo, B. T., Robinson, C. V., and Bowers, M. T. (2009) Amyloid-β 2 protein oligomerization and the importance of tetramers and dodecamers in the aetiology of Alzheimer’s disease. Nat. Chem. 1, 326–331

16. Walsh, D. M., Hartley, D. M., Kusumoto, Y., Fezoui, Y., Condron, M. M., Lomakin, A., Benedek, G. B., Selkoe, D. J., and Teplow, D. B. (1999) Amyloid β-protein fibrillogenesis. Structure and biological activity of protofibrillar intermediates. J. Biol. Chem. 10.1074/jbc.274.36.25945

17. Fändrich, M. (2012) Oligomeric intermediates in amyloid formation: Structure determination and mechanisms of toxicity. J. Mol. Biol. 10.1016/j.jmb.2012.01.006

18. Biancalana, M., and Koide, S. (2010) Molecular mechanism of Thioflavin-T binding to amyloid fibrils. Biochim. Biophys. Acta - Proteins Proteomics. 1804, 1405–1412

19. Tiiman, A., Jarvet, J., Gräslund, A., and Vukojevic, V. (2015) Heterogeneity and intermediates turnover during amyloid-β (Aβ) peptide aggregation studied by Fluorescence Correlation Spectroscopy. Biochemistry. 10.1021/acs.biochem.5b00976

20. Michaels, T. C. T., Šarić, A., Curk, S., Bernfur, K., Arosio, P., Meisl, G., Dear, A. J., Cohen, S. I. A., Vendruscolo, M., Dobson, C. M., Linse, S., and Knowles, T. P. J. (2020) Dynamics of oligomer populations formed during the aggregation of Alzheimer’s Aβ42 peptide. bioRxiv. 10.1101/2020.01.08.897488

21. Linse, S. (2017) Monomer-dependent secondary nucleation in amyloid formation. Biophys. Rev. 9, 329–338

22. Österlund, N., Moons, R., Ilag, L. L., Sobott, F., and Gräslund, A. (2019) Native Ion Mobility-Mass Spectrometry Reveals the Formation of β-Barrel Shaped Amyloid-β Hexamers in a Membrane-Mimicking Environment. J. Am. Chem. Soc. 141, 10440–10450

23. Hageman, J., Rujano, M. A., van Waarde, M. A. W. H., Kakkar, V., Dirks, R. P., Govorukhina, N., Oosterveld-Hut, H. M. J., Lubsen, N. H., and Kampinga, H. H. (2010) A DNAJB Chaperone Subfamily with HDAC-Dependent Activities Suppresses Toxic Protein Aggregation. Mol. Cell. 10.1016/j.molcel.2010.01.001

24. Månsson, C., Arosio, P., Hussein, R., Kampinga, H. H., Hashem, R. M., Boelens, W. C., Dobson, C. M., Knowles, T. P. J., Linse, S., and Emanuelsson, C. (2014) Interaction of the molecular chaperone DNAJB6 with growing amyloid-beta 42 (Aβ42) aggregates leads to sub-stoichiometric inhibition of amyloid formation. J. Biol. Chem. 289, 31066–31076

25. Månsson, C., Kakkar, V., Monsellier, E., Sourigues, Y., Härmark, J., Kampinga, H. H., Melki, R., and Emanuelsson, C. (2014) DNAJB6 is a peptide-binding chaperone which can suppress amyloid fibrillation of polyglutamine peptides at substoichiometric molar ratios. Cell Stress Chaperones. 19, 227–239

26. Kakkar, V., Månsson, C., de Mattos, E. P., Bergink, S., van der Zwaag, M., van Waarde, M. A. W. H., Kloosterhuis, N. J., Melki, R., van Cruchten, R. T. P., Al-Karadaghi, S., Arosio, P., Dobson, C. M., Knowles, T. P. J., Bates, G. P., van Deursen, J. M., Linse, S., van de Sluis, B., Emanuelsson, C., and Kampinga, H. H. (2016) The S/T-Rich Motif in the DNAJB6 Chaperone Delays Polyglutamine Aggregation and the Onset of Disease in a Mouse Model. Mol. Cell. 62, 272–283

27. Hunter, P. J., Swanson, B. J., Haendel, M. A., Lyons, G. E., and Cross, J. C. (1999) Mrj encodes a DnaJ-related co-chaperone that is essential for murine placental development. Development. 126, 1247–1258

28. Månsson, C., Van Cruchten, R. T. P., Weininger, U., Yang, X., Cukalevski, R., Arosio, P., Dobson, C. M., Knowles, T., Akke, M., Linse, S., and Emanuelsson, C. (2018) Conserved S/T Residues of the Human Chaperone DNAJB6 Are Required for Effective Inhibition of Aβ42 Amyloid Fibril Formation. Biochemistry. 57, 4891–4892

29. Söderberg, C. A. G., Månsson, C., Bernfur, K., Rutsdottir, G., Härmark, J., Rajan, S., Al-Karadaghi, S., Rasmussen, M., Höjrup, P., Hebert, H., and Emanuelsson, C. (2018) Structural modelling of the DNAJB6 oligomeric chaperone shows a peptide-binding cleft lined with conserved S/T-residues at the dimer interface. Sci. Rep. 10.1038/s41598-018-23035-9

30. Testa, L., Brocca, S., and Grandori, R. (2011) Charge-surface correlation in electrospray ionization of folded and unfolded proteins. Anal. Chem. 83, 6459–6463

31. Liu, L., Dong, X., Liu, Y., Österlund, N., Gräslund, A., Carloni, P., and Li, J. (2019) Role of hydrophobic residues for the gaseous formation of helical motifs. Chem. Commun. 10.1039/c9cc01898k

32. Pagel, K., Natan, E., Hall, Z., Fersht, A. R., and Robinson, C. V. (2013) Intrinsically disordered p53 and its complexes populate compact conformations in the gas phase. Angew. Chemie - Int. Ed. 10.1002/anie.201203047

33. Marsh, J. A., and Teichmann, S. A. (2011) Relative solvent accessible surface area predicts protein conformational changes upon binding. Structure. 19, 859–867

34. Li, J., Santambrogio, C., Brocca, S., Rossetti, G., Carloni, P., and Grandori, R. (2016) Conformational effects in protein electrospray-ionization mass spectrometry. Mass Spectrom. Rev. 35, 111–122

35. Hoyer, W., Gronwall, C., Jonsson, A., Stahl, S., and Hard, T. (2008) Stabilization of a beta-hairpin in monomeric Alzheimer’s amyloid-beta peptide inhibits amyloid formation. Proc Natl Acad Sci U S A. 105, 5099–5104

36. Lindberg, H., Härd, T., Löfblom, J., and Ståhl, S. (2015) A truncated and dimeric format of an Affibody library on bacteria enables FACS-mediated isolation of amyloid-beta aggregation inhibitors with subnanomolar affinity. Biotechnol. J. 10.1002/biot.201500131

37. Cheon, M., Chang, I., Mohanty, S., Luheshi, L. M., Dobson, C. M., Vendruscolo, M., and Favrin, G. (2007) Structural reorganisation and potential toxicity of oligomeric species formed during the assembly of amyloid fibrils. PLoS Comput. Biol. 10.1371/journal.pcbi.0030173

38. Ghadami, S. A., Chia, S., Ruggeri, F. S., Meisl, G., Bemporad, F., Habchi, J., Cascella, R., Dobson, C. M., Vendruscolo, M., Knowles, T. P. J., and Chiti, F. (2020) Transthyretin inhibits primary and secondary nucleation of amyloid-β peptide aggregation and reduces the toxicity of its oligomers. Biomacromolecules. 10.1021/acs.biomac.9b01475

39. Yokoyama, T., Mizuguchi, M., Nabeshima, Y., Kusaka, K., Yamada, T., Hosoya, T., Ohhara, T., Kurihara, K., Tomoyori, K., Tanaka, I., and Niimura, N. (2012) Hydrogen-bond network and pH sensitivity in transthyretin: Neutron crystal structure of human transthyretin. J. Struct. Biol. 10.1016/j.jsb.2011.12.022

40. Testa, L., Brocca, S., Santambrogio, C., D’Urzo, A., Habchi, J., Longhi, S., Uversky, V. N., and Grandori, R. (2013) Extracting structural information from charge-state distributions of intrinsically disordered proteins by non-denaturing electrospray-ionization mass spectrometry. Intrinsically Disord. Proteins. 10.4161/idp.25068

41. Frimpong, A. K., Abzalimov, R. R., Uversky, V. N., and Kaltashov, I. A. (2010) Characterizationof intrinsically disordered proteins with electrospray inization mass spectrometry: Conformationl heterogeneity of α-synuciein. Proteins Struct. Funct. Bioinforma. 10.1002/prot.22604

42. Konijnenberg, A., Ranica, S., Narkiewicz, J., Legname, G., Grandori, R., Sobott, F., and Natalello, A. (2016) Opposite Structural Effects of Epigallocatechin-3-gallate and Dopamine Binding to α-Synuclein. Anal. Chem. 10.1021/acs.analchem.6b00731

43. Walsh, D. M., Thulin, E., Minogue, A. M., Gustavsson, N., Pang, E., Teplow, D. B., and Linse, S. (2009) A facile method for expression and purification of the Alzheimer’s disease-associated amyloid β-peptide. FEBS J. 276, 1266–1281

